# Pre-stress functional brain connectivity is associated with subsequent anxiety during war-related stress

**DOI:** 10.64898/2026.10.24.761100

**Authors:** N. Boshnak, N. Danan, H. Keren

**Author notes:** These authors contributed equally to this work (N. Boshnak, N. Danan).

## Abstract

Individuals exposed to major real-world stressors show markedly different psychological outcomes, but whether this heterogeneity is associated with pre-existing neural characteristics remains unclear because neuroimaging is rarely performed beforehand. We acquired resting-state fMRI in healthy adults before the unforeseen onset of war-related stress. Approximately 13 months later, during ongoing war-related stress, participants (n = 31) were assessed for anxiety, depressive symptoms, perceived stress and post-traumatic stress symptoms. Among 144 associations spanning 36 predefined corticolimbic and reward-related connections and four outcomes, lower baseline functional connectivity between the right nucleus accumbens and right amygdala was associated with greater subsequent anxiety. This association survived permutation-based family-wise error correction and adjustment for baseline depressive symptoms, age and sex. These findings provide preliminary prospective evidence that pre-existing functional organization of threat- and reward-related circuitry is associated with individual differences in anxiety during prolonged adversity.

## Introduction

Prolonged real-world stress is a well-established risk factor for the development of anxiety, depression, and trauma-related symptoms^1,2^. Yet individuals exposed to comparable stressors often follow markedly different psychological trajectories, ranging from minimal symptoms to persistent anxiety, depression, or trauma-related symptoms^3,4^. The neurobiological factors underlying this heterogeneity remain incompletely understood, limiting our ability to identify individuals at greater risk of adverse psychological outcomes following adversity. Understanding the neurobiological basis of these individual differences therefore remains a central challenge in stress neuroscience.

A major obstacle to addressing this question is that neuroimaging is typically performed after exposure to major stressors or after psychological symptoms have already emerged^5^. Consequently, many studies cannot readily determine whether observed neural characteristics were present before adversity or emerged in response to it. Prospective neuroimaging performed before the onset of such stressors provides a valuable opportunity to clarify the temporal relationship between brain organization and psychological responses to adversity and to distinguish between these possibilities.

Recent prospective neuroimaging studies have begun to demonstrate that neural characteristics measured before the onset of major stressors are associated with subsequent psychological outcomes. For example, Zhang et al.^6^ showed that acute stress-induced reconfiguration of large-scale brain networks, assessed before trauma exposure, prospectively predicted subsequent stress-related symptoms in police recruits. More recently, longitudinal work conducted around the COVID-19 pandemic demonstrated that pre-stress functional connectivity of the amygdala and nucleus accumbens with other brain regions was associated with subsequent perceived stress and anxiety following a real-world stressor^7^. Together, these studies highlight the relevance of neural characteristics measured before adversity for understanding individual differences in subsequent mental health. Importantly, previous work has primarily examined neural responses to experimentally evoked acute stress or neural connectivity of the amygdala and nucleus accumbens considered separately. However, whether intrinsic resting-state functional connectivity between these regions, measured before the onset of prolonged real-world stress in the absence of experimental manipulation, is associated with subsequent psychological outcomes remains unclear.

Among the neural systems implicated in stress adaptation, interactions between threat-and reward-related circuits have been shown to be particularly important. The amygdala plays a central role in evaluating biologically salient and potentially threatening information^8^, whereas the nucleus accumbens contributes to reward processing, motivation, and goal-directed behaviour^9^. Functional interactions between these regions may therefore support the integration of threat and motivational information required to balance defensive responding with continued engagement in rewarding and adaptive behaviour under conditions of uncertainty. Consistent with this possibility, preclinical evidence implicates amygdala–nucleus accumbens circuitry in behavioural responses to chronic stress^10^.

Here, we investigated whether intrinsic functional brain organization measured before the unforeseen onset of war-related stress was prospectively associated with subsequent psychological outcomes. Resting-state fMRI was acquired in psychologically healthy adults approximately 13 months before follow-up and before the onset of war-related stress. At follow-up, during ongoing war-related stress, we assessed anxiety, depressive symptoms, perceived stress, and post-traumatic stress symptoms in the same individuals. We examined whether pre-stress functional connectivity across 36 predefined corticolimbic and reward-related connections was associated with individual differences in these subsequent psychological outcomes.

## Results

The study design and assessment timeline are summarized in **Fig. 1A**. At baseline (T1), resting-state functional MRI data were acquired and in-scanner mood ratings were collected in 31 participants. Clinical follow-up assessments (T2) were conducted a mean of 13.2 months after baseline (SD = 2.4; range, 8.5–16.1 months) during ongoing war-related stress and included measures of depressive symptoms (CES-D), anxiety symptoms (GAD-7), perceived stress (PSS), and post-traumatic stress symptoms (PCL-5). At T2, participants also reported their war-related exposure, including reserve service, war-related harm, relocation, alarm exposure, and socioeconomic deterioration. Specifically, 15 participants (48.4%) reported that they or someone close to them was serving in the military reserves, 2 (6.5%) reported war-related harm to themselves or an immediate family member, 9 (29.0%) reported relocating from their home, 19 (61.3%) reported exposure to alarms once per day or more, and 11 (35.5%) reported socioeconomic deterioration. War-related exposure characteristics are summarized in **Supplementary Table S3**.

**Figure 1.**
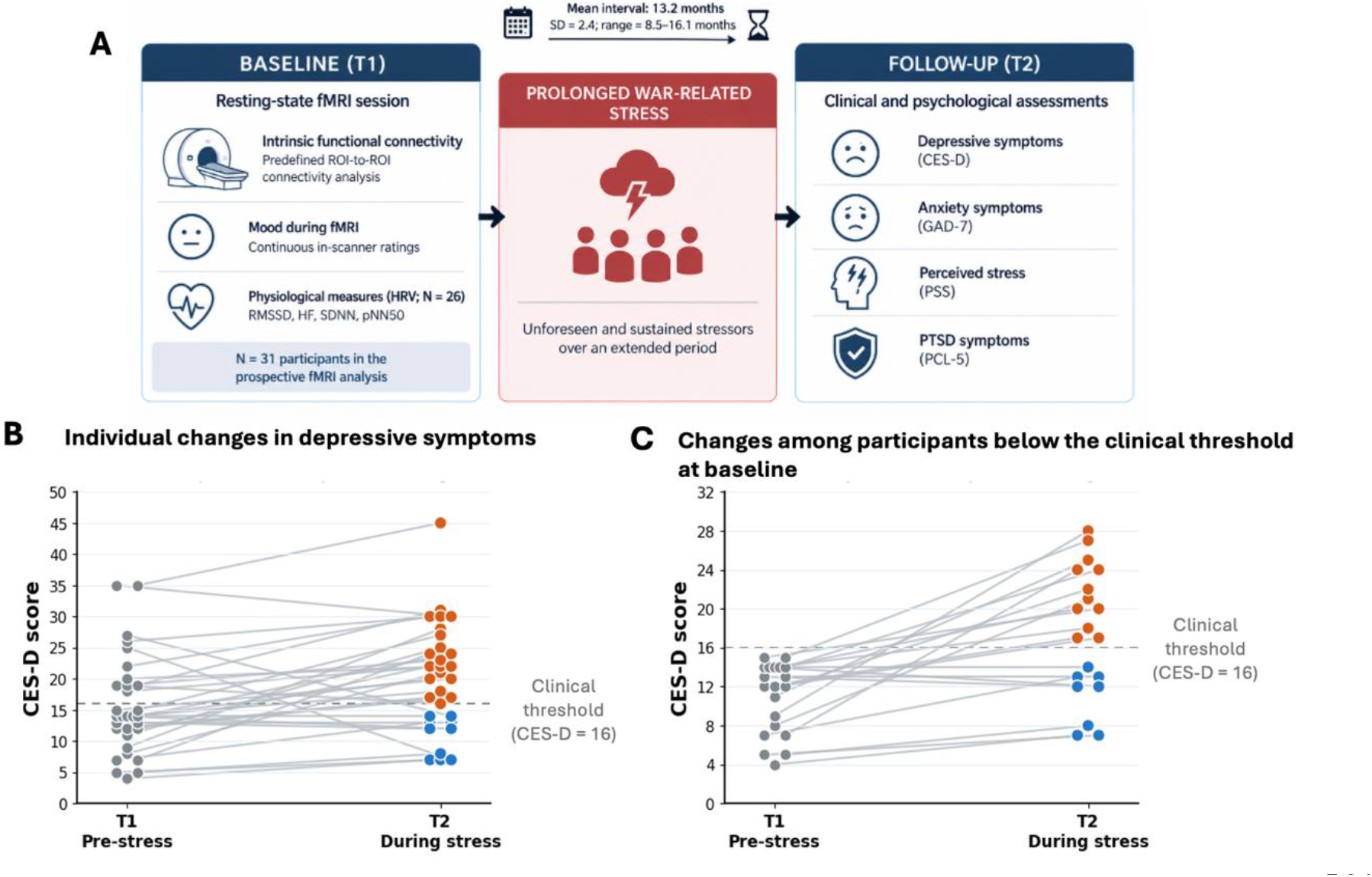
Study design and prospective assessment during ongoing war-related stress. **(A)** Study overview. Resting-state functional MRI was acquired at baseline (T1) in the prospective fMRI sample (n = 31). Clinical follow-up assessments were conducted a mean of 13.2 months later (SD = 2.4; range, 8.5–16.1 months), during ongoing war-related stress (T2). **(B)** Individual changes in depressive symptoms (CES-D) from baseline (T1) to follow-up (T2). Each line represents one participant. **(C)** Changes in depressive symptoms among participants who scored below the conventional clinical threshold at baseline (CES-D <16; n = 20), demonstrating substantial inter-individual variability in depressive symptom severity at follow-up. In **B** and **C**, the dashed line indicates the conventional clinical threshold for elevated depressive symptoms (CES-D ≥16). Baseline observations are shown in gray; follow-up observations are shown in orange for participants with CES-D ≥16 and in blue for participants with CES-D <16.

Psychological outcomes at follow-up varied substantially across participants. CES-D scores ranged from 7 to 45 (mean ± SD, 20.0 ± 8.6), GAD-7 scores from 0 to 15 (5.4 ± 3.6), PSS scores from 16 to 37 (27.1 ± 5.7), and PCL-5 scores from 1 to 61 (22.5 ± 15.6). Follow-up questionnaire scores did not differ significantly between male and female participants after FDR correction (Supplementary Fig. S2). Depressive symptom trajectories ranged from persistently low symptom levels to substantial increases from baseline to follow-up (**Fig. 1B**). At baseline, 11 of 31 participants (35.5%) exceeded the conventional clinical threshold for elevated depressive symptoms (CES-D ≥ 16). Of these participants, 9 remained above the threshold at follow-up, whereas 2 scored below it. At follow-up, 21 of 31 participants (67.7%) exceeded the clinical threshold, whereas 10 scored below it. **Figure 1C** further illustrates this inter-individual heterogeneity among the 20 participants who scored below the clinical threshold at baseline (CES-D < 16). At follow-up, CES-D scores in this subgroup ranged from 7 to 28: 12 participants exceeded the clinical threshold, whereas 8 remained below it. Thus, participants without elevated depressive symptoms at baseline exhibited markedly divergent depressive symptom trajectories during ongoing war-related stress.

Participants with elevated depressive symptoms at follow-up also exhibited higher levels of anxiety symptoms, perceived stress, and post-traumatic stress symptoms, together with lower interoceptive awareness (**Supplementary Fig. S3**).

### Right nucleus accumbens–amygdala connectivity identified as a pre-stress neural correlate of subsequent anxiety symptoms

To identify neural circuits associated with inter-individual differences in psychological outcomes, we examined associations between baseline resting-state functional connectivity among nine predefined regions of interest (36 ROI-to-ROI functional connections; **Fig. 2B**) and four follow-up psychological outcomes (CES-D, GAD-7, PSS, and PCL-5).

**Figure 2.**
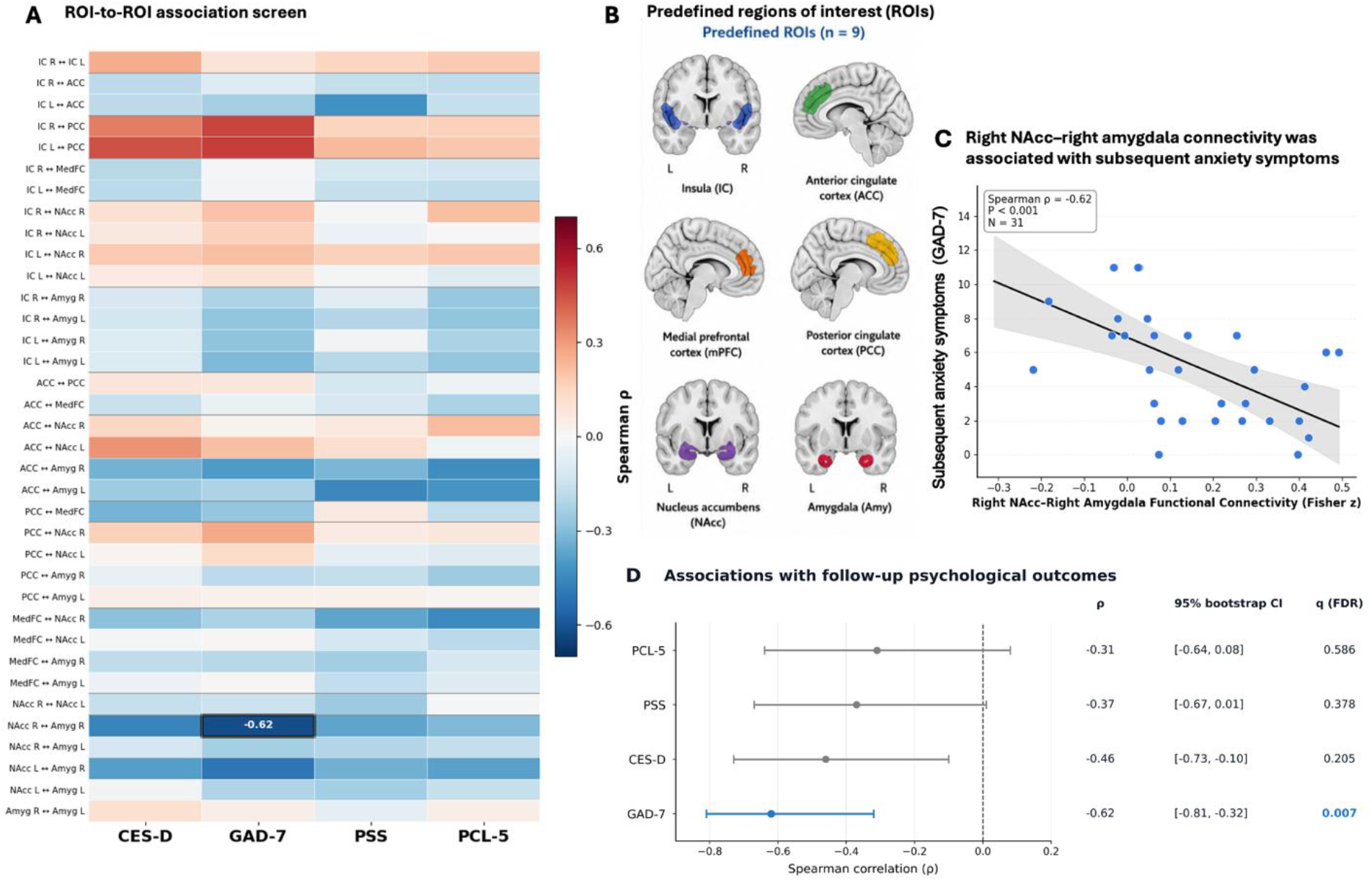
ROI-to-ROI connectivity screening identifies an association between baseline right nucleus accumbens–right amygdala connectivity and subsequent anxiety symptoms. **(A)** Pairwise association screen relating baseline resting-state functional connectivity across 36 predefined ROI-to-ROI connections to four follow-up psychological outcomes: depressive symptoms (CES-D), anxiety symptoms (GAD-7), perceived stress (PSS), and post-traumatic stress symptoms (PCL-5). Colors indicate Spearman correlation coefficients (ρ). For each psychological outcome, Benjamini–Hochberg false discovery rate (FDR) correction was applied across the 36 predefined functional connections. The black square denotes the association between baseline right NAcc–right amygdala connectivity and follow-up GAD-7 scores, which remained significant after global Benjamini–Hochberg FDR correction across all 144 connectivity–outcome associations and after permutation-based maximum-statistic correction. **(B)** Anatomical locations of the nine predefined regions of interest (ROIs) included in the ROI-to-ROI functional connectivity analysis: bilateral insula (IC), anterior cingulate cortex (ACC), medial prefrontal cortex (mPFC), posterior cingulate cortex (PCC), bilateral nucleus accumbens (NAcc), and bilateral amygdala (Amy). **(C)** Association between baseline right NAcc–right amygdala functional connectivity and follow-up anxiety symptoms (GAD-7), with individual participant values shown. Each point represents one participant. The solid line indicates the linear regression fit, and the shaded area denotes its 95% confidence interval. The reported association was assessed using Spearman’s rank correlation. **(D)** Associations between baseline right NAcc–right amygdala functional connectivity and each follow-up psychological outcome. Horizontal lines indicate 95% bootstrap confidence intervals estimated from 10,000 resamples. FDR-adjusted q values from the corresponding outcome-specific analyses across 36 connections are shown.

This screening revealed substantial variability in both the strength and direction of associations across the examined connections (**Fig. 2A**). Among the 144 connectivity–outcome associations tested, baseline connectivity between the right nucleus accumbens and the right amygdala was strongly associated with subsequent anxiety symptoms (Spearman’s ρ = −0.619, permutation P = 3.2 × 10^-4^; **Fig. 2C**). This association remained significant after Benjamini–Hochberg FDR correction across all 144 connectivity–outcome associations (q = 0.046), as well as after permutation-based maximum-statistic correction controlling the family-wise error rate across the same set of tests (P_FWE = 0.032). It also remained significant after outcome-specific FDR correction across the 36 connections examined in relation to anxiety symptoms (q = 0.007). No other connectivity–outcome association remained significant after either global correction.

For the identified right nucleus accumbens–right amygdala connection, weaker negative associations were observed with the three other psychological outcomes-depressive symptoms (CES-D), perceived stress (PSS), and post-traumatic stress symptoms (PCL-5)-none of which remained significant after outcome-specific FDR correction (**Fig. 2D**).

Together, these findings identify right nucleus accumbens–right amygdala connectivity as a prospective neural correlate of subsequent anxiety symptoms within the examined neural network.

To examine whether baseline connectivity was associated with participants’ concurrent affective state or within-scan mood dynamics, we tested its associations with six in-scanner mood measures: the first, mean, and final mood ratings, mood variability, mood slope, and overall mood change. Right nucleus accumbens–right amygdala connectivity was not significantly associated with any of these measures after FDR correction (**Supplementary Fig. S4**).

### Baseline right nucleus accumbens–right amygdala connectivity is associated with subsequent anxiety symptoms beyond baseline depressive symptoms

Having identified baseline right nucleus accumbens–right amygdala connectivity as a strong neural correlate of subsequent anxiety symptoms, we next examined whether this association remained evident after accounting for baseline depressive symptom severity. Baseline depressive symptoms were positively associated with subsequent anxiety symptoms (Spearman’s ρ = 0.43, P = 0.017; Fig. 3A).

**Figure 3.**
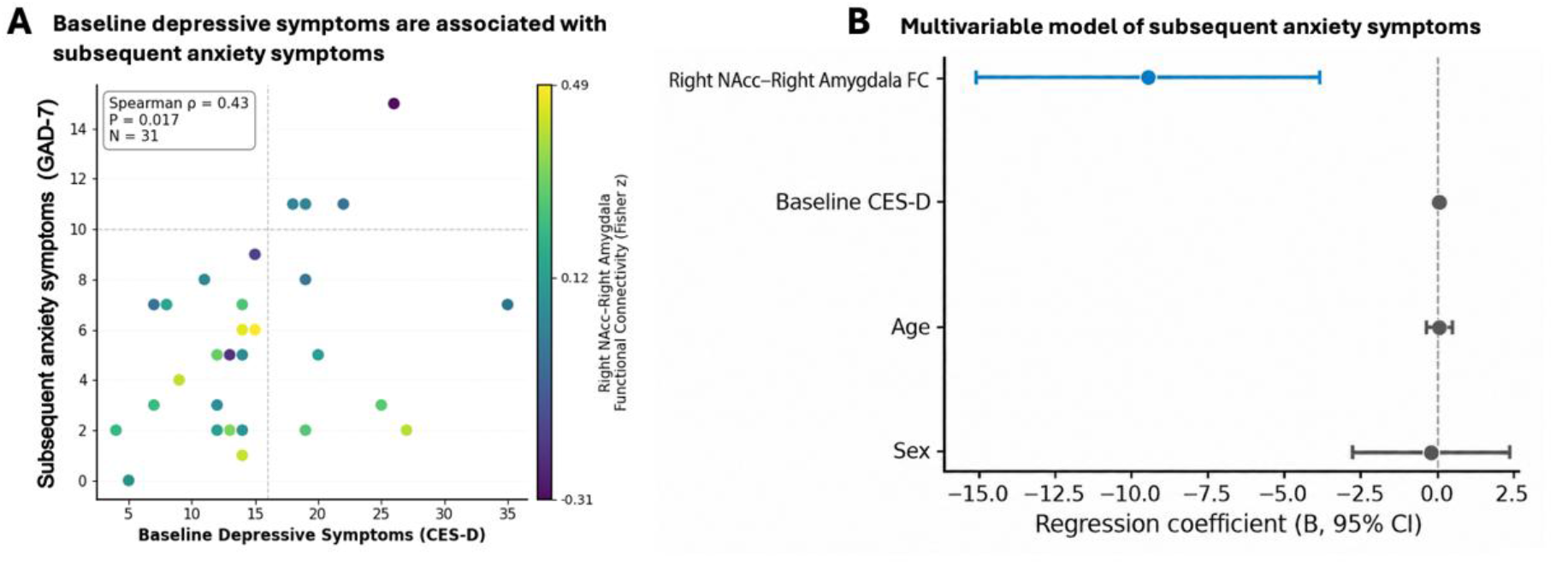
Baseline right nucleus accumbens–right amygdala connectivity is associated with subsequent anxiety symptoms after adjustment for baseline depressive symptoms, age, and sex. **(A)** Association between baseline depressive symptoms (CES-D) and subsequent anxiety symptoms (GAD-7). Point color represents baseline right nucleus accumbens–right amygdala functional connectivity, illustrating variability in circuit connectivity among participants with comparable baseline depressive symptom severity. **(B)** Unstandardized regression coefficients (B) and 95% confidence intervals from the multivariable model with subsequent anxiety symptoms as the dependent variable and baseline right nucleus accumbens–right amygdala functional connectivity, baseline CES-D score, age, and sex as predictors. The dashed vertical line indicates zero. Full coefficient estimates, standardized coefficients, confidence intervals, P values, and model fit statistics are reported in **Supplementary Table S1**.

We fitted a multivariable regression model that included baseline right nucleus accumbens–right amygdala connectivity as the predictor of interest and baseline CES-D score, age, and sex as covariates. Lower baseline connectivity remained significantly associated with greater subsequent anxiety symptoms after adjustment for baseline depressive symptoms, age, and sex (B = −9.50, standardized β = −0.53, 95% CI [−15.14, −3.86], P = 0.002; **Fig. 3B** and **Supplementary Table S1**). Baseline depressive symptoms showed a positive association that did not reach statistical significance after adjustment (P = 0.094), whereas neither age nor sex was significantly associated with subsequent anxiety symptoms. The overall model explained 44.3% of the variance in subsequent anxiety symptoms (adjusted R² = 0.357; model P = 0.003).

### The connectivity–anxiety association remains stable across complementary sensitivity analyses

Given the modest sample size and concerns regarding the reliability of brain–behaviour associations in small samples, we conducted a series of complementary sensitivity analyses to evaluate the stability of the primary association between right nucleus accumbens–right amygdala connectivity and subsequent anxiety symptoms (**Supplementary Table S2** and **Supplementary Fig. S5**). Leave-one-out analysis showed that the correlation remained stable across all iterations (ρ range, −0.668 to −0.580), indicating that no single participant disproportionately influenced the observed association. The 95% bootstrap confidence interval, estimated from 10,000 resamples, excluded zero (95% CI, −0.807 to −0.314). A nonparametric permutation test based on 100,000 permutations yielded a comparable result (P = 3.2 × 10^-4^). The association also remained significant under a permutation-based maximum-statistic correction controlling the family-wise error rate across the 36 predefined functional connections examined in relation to anxiety symptoms (P_FWE = 0.009).

We next examined whether the observed association between baseline connectivity and subsequent anxiety remained evident after accounting for measured differences in war-related exposure, including reserve service, war-related harm, relocation, alarm exposure, and socioeconomic deterioration (**Supplementary Tables S4–S5**). To examine this, we created a security-exposure index comprising reserve service, war-related harm, relocation, and alarm exposure once per day or more (range, 0–4), as well as a full exposure index that additionally incorporated socioeconomic deterioration (range, 0–5). Neither index was associated with baseline right nucleus accumbens–right amygdala connectivity (all |ρ| ≤ 0.14, all P ≥ 0.44). In uncorrected secondary analyses, both the security-exposure index (ρ = 0.38, P = 0.037) and the full exposure index (ρ = 0.44, P = 0.012) were positively associated with subsequent anxiety symptoms. When examined separately, socioeconomic deterioration was likewise positively associated with subsequent anxiety symptoms (ρ = 0.42, P = 0.018), but not with baseline connectivity (ρ = −0.22, P = 0.237; **Supplementary Table S4**). Importantly, the estimated connectivity coefficient remained similar in magnitude (B range, −9.50 to −8.20) and statistically significant (all P ≤ 0.009) across models incrementally adjusting for age, sex, and the war-related exposure measures, including a model jointly adjusting for the security-exposure index and the socioeconomic-deterioration indicator (**Supplementary Table S5** and **Supplementary Fig. S6**). Together, these analyses indicate that the association between baseline connectivity and subsequent anxiety remained evident after adjustment for measured individual differences in war-related exposure.

## DISCUSSION

A fundamental challenge in stress neuroscience is understanding why individuals exposed to prolonged adversity develop markedly different psychological outcomes. Here, leveraging a prospective design, we examined intrinsic functional brain organization before the unforeseen onset of war-related stress, approximately 13 months before follow-up, and related it to subsequent psychological outcomes assessed during ongoing war-related stress. Among 144 connectivity–outcome associations, baseline functional connectivity between the right nucleus accumbens and the right amygdala showed a strong association with subsequent anxiety symptoms that survived global correction for multiple comparisons. Lower baseline connectivity was associated with greater subsequent anxiety, and this association remained significant after adjustment for baseline depressive symptoms, age and sex. It also remained evident in models adjusting for measured differences in war-related exposure.

Moreover, this baseline connectivity was not significantly associated with concurrent in-scanner mood level or dynamics after FDR correction, suggesting that the prospective association was not simply attributable to participants’ momentary affective state at the time of scanning. Overall, these findings identify intrinsic amygdala–nucleus accumbens connectivity as a prospective neural correlate of subsequent anxiety severity during prolonged real-world adversity.

Our findings complement previous prospective research showing that brain characteristics measured before the onset of major stressors are associated with subsequent psychological outcomes^5–7^. In adolescents studied before the COVID-19 pandemic, pre-pandemic nucleus accumbens connectivity with other brain regions was associated with later anxiety^7^. The present study identifies an association involving pairwise nucleus accumbens–amygdala functional connectivity in adults exposed to a distinct form of prolonged adversity. These studies thus provide complementary evidence implicating baseline reward-related circuitry in subsequent anxiety, while our study extends these findings to a different context of real-world adversity and additionally implicates the amygdala as part of this circuitry.

The amygdala–nucleus accumbens circuit is well positioned to support adaptation to prolonged stress through the integration of threat-related and motivational information. Functional interactions between these regions may contribute to balancing defensive responses with reward engagement, goal-directed behaviour, and behavioural flexibility as environmental demands change. Human neuroimaging research has also implicated nucleus accumbens–amygdala functional connectivity in avoidance learning^11^. These observations are consistent with contemporary models in which stress adaptation emerges from coordinated interactions among corticolimbic and reward-related circuits rather than from isolated brain regions^1,2,12^. Within this framework, individual variation in intrinsic amygdala–nucleus accumbens connectivity may reflect differences in the functional organization of neural systems involved in responding to prolonged adversity.

Preclinical research provides a mechanistic framework for interpreting the potential relevance of this circuitry. In mice, chronic social stress altered basal amygdala–nucleus accumbens pathway activity during reward learning and reward-to-effort valuation. Experimental inhibition or activation of this pathway reproduced distinct stress-related impairments in reward behaviour^10^. These findings provide causal evidence that amygdala–nucleus accumbens pathway function contributes to behavioural responses to chronic stress. Although human resting-state functional connectivity cannot establish causal or projection-specific mechanisms, the present prospective association is consistent with experimental evidence implicating interactions between threat- and reward-related systems in responses to adversity and provides a translational link between mechanistic circuit neuroscience and individual differences in psychological outcomes during prolonged real-world stress.

An important consideration is whether the observed association reflects individual differences in exposure to war-related stressors rather than pre-existing functional brain organization. Although participants shared the broader context of an ongoing regional conflict, they differed in their exposure to reserve service, war-related harm, relocation, alarm exposure, and socioeconomic deterioration. In uncorrected secondary analyses, greater cumulative war-related exposure and socioeconomic deterioration were associated with greater subsequent anxiety. However, neither cumulative exposure index nor socioeconomic deterioration was associated with baseline right nucleus accumbens–right amygdala connectivity. Moreover, the connectivity–anxiety association remained evident after adjustment for the measured exposure indicators, including a model jointly accounting for the security-exposure index and the socioeconomic-deterioration indicator. These findings suggest that, although measured war-related exposure was associated with anxiety severity, it did not readily account for the association between baseline connectivity and subsequent anxiety.

The strongest prospective association observed in the present study involved anxiety symptoms, whereas associations with depressive symptoms, perceived stress and post-traumatic stress symptoms were weaker and did not survive correction for multiple comparisons. However, these findings should not necessarily be interpreted as evidence for an anxiety-specific circuit. The psychological outcomes were substantially intercorrelated (**Supplementary Fig. S1**), consistent with dimensional models of psychopathology in which anxiety, depression and trauma-related symptoms represent overlapping but partially dissociable dimensions of psychological distress^13^. The observed pattern may reflect differences in effect magnitude or symptom expression across domains, together with limited statistical power to detect more modest associations. Furthermore, a statistically significant association with anxiety and nonsignificant associations with other outcomes do not establish that these associations differ significantly from one another.

The right-lateralized nature of the observed association warrants further investigation. Previous research suggests functional asymmetries in amygdala-mediated emotional processing, although findings vary across experimental paradigms^8,14^. However, because we did not directly test whether the right-sided association was stronger than the corresponding left-sided associations, this pattern should be considered preliminary. Several limitations should be acknowledged. First, anxiety symptoms were not assessed before the onset of war-related stress. We therefore cannot determine whether the observed association reflects pre-existing individual differences in anxiety, differential changes in anxiety during prolonged adversity, or both. Although baseline depressive symptoms were measured and included in the adjusted analyses, and depressive and anxiety symptoms were strongly correlated at follow-up, baseline depressive symptoms cannot substitute for a baseline anxiety assessment. Second, the relatively small sample warrants cautious interpretation of the magnitude and stability of the observed association. Although the connectivity–anxiety relationship survived stringent correction across 144 examined associations and remained relatively stable across complementary robustness analyses, independent replication in larger samples is needed to establish its reliability and generalizability. Third, functional connectivity is an indirect correlational measure and cannot establish the directionality or causal mechanisms of amygdala–nucleus accumbens interactions. In addition, functional connectivity was assessed at a single time point and thus the study cannot determine whether the observed pattern reflects a stable individual characteristic or a temporally varying neural state. Fourth, war-related exposure was assessed at follow-up using a limited set of binary indicators. Although these measures captured several salient sources of exposure, they may not fully reflect the timing, severity, duration, or subjective experience of adversity. Residual confounding by unmeasured aspects of war-related exposure therefore cannot be excluded. Finally, attrition differed by sex: female participants were more likely than male participants to decline follow-up participation, resulting in their underrepresentation in the final sample. Although sex was included as a covariate, this adjustment cannot rule out selection bias arising from differential attrition, which may limit the generalizability of the findings.

In conclusion, intrinsic functional connectivity between the right nucleus accumbens and the right amygdala, measured before the unforeseen onset of war-related stress and approximately 13 months before follow-up, was prospectively associated with subsequent anxiety symptoms. These findings suggest that variation in functional connectivity between the amygdala and nucleus accumbens is associated with individual differences in anxiety during prolonged real-world adversity. Together with previous prospective observations and experimental evidence implicating amygdala–nucleus accumbens circuitry in responses to chronic stress, the present findings highlight the potential relevance of specific features of pre-existing functional brain organization for understanding individual differences in psychological outcomes following major adversity.

## METHODS

### Participants and prospective study design

Participants were drawn from a longitudinal neuroimaging cohort^15^ originally recruited from universities and the surrounding community. Participants were screened before **enrolment**. Exclusion criteria included standard MRI contraindications (e.g., metallic implants or pacemakers), claustrophobia, epilepsy, pregnancy, medical conditions that could interfere with MRI scanning, a self-reported history of neurological or psychiatric disorders, neurodevelopmental conditions or cognitive impairment, and current use of medications with potential effects on the central nervous system.

The present prospective study comprised two assessment time points. At baseline (T1), participants underwent resting-state functional MRI interleaved with repeated in-scanner mood assessments and an assessment of depressive symptoms. Forty-four participants had usable baseline resting-state fMRI data (mean age ± SD = 22.6 ± 3.1 years; range, 18–31 years; 23 male and 21 female). At a mean of 13.2 months after baseline (SD = 2.4; range, 8.5–16.1 months), following the unforeseen onset of war-related stress, all 44 participants were invited to complete a follow-up assessment during ongoing war-related stress (T2), including measures of depressive symptoms, anxiety symptoms, perceived stress, post-traumatic stress symptoms, and interoceptive awareness. Of these, 13 declined to participate in the follow-up assessment. Participants who completed the follow-up assessment did not differ from those who declined in age (22.4 ± 2.9 vs. 23.3 ± 3.5 years; Welch’s t = −0.86, P = 0.399) or baseline depressive symptoms (CES-D: 15.6 ± 7.9 vs. 17.2 ± 7.7; Welch’s t = −0.62, P = 0.544). However, female participants were more likely than male participants to decline participation (11 of 21 women vs. 2 of 23 men; Fisher’s exact test, P = 0.002). The final sample for the prospective analyses therefore comprised 31 participants with both usable baseline resting-state fMRI data and complete follow-up assessments (mean age ± SD = 22.4 ± 2.9 years; range, 18–31 years; 21 male and 10 female). Demographic and clinical characteristics of the final sample are summarized in **Table 1**.

**Table 1.** Demographic and clinical characteristics of the fMRI cohort. Data are presented as mean ± SD unless otherwise indicated. Elevated depressive symptoms were defined as a CES-D score ≥16 at follow-up (T2). CES-D, Center for Epidemiologic Studies Depression Scale; GAD-7, Generalized Anxiety Disorder-7; PSS, Perceived Stress Scale; PCL-5, PTSD Checklist for DSM-5; MAIA-2, Multidimensional Assessment of Interoceptive Awareness, Version 2.

| Characteristic | Total (N = 31) |
| --- | --- |
| Age, years | 22.4 ± 2.9 |
| Age range | 18–31 |
| **Sex, n (%)** |  |
| Male | 21 (67.7%) |
| Female | 10 (32.3%) |
| CES-D (T1) | 15.6 ± 7.9 |
| CES-D (T2) | 20.0 ± 8.6 |
| Elevated depressive symptoms at T2, n (%) | 21 (67.7%) |
| GAD-7 | 5.4 ± 3.6 |
| PSS | 27.1 ± 5.7 |
| PCL-5 | 22.5 ± 15.6 |
| MAIA-2 total score | 4.0 ± 0.6 |

Values are presented as mean ± SD unless otherwise indicated. Elevated depressive symptoms were defined as a CES-D score ≥16 at follow-up (T2). CES-D, Center for Epidemiologic Studies Depression Scale; GAD-7, Generalized Anxiety Disorder-7; PSS, Perceived Stress Scale; PCL-5, PTSD Checklist for DSM-5; MAIA-2, Multidimensional Assessment of Interoceptive Awareness, Version 2.

All participants provided written informed consent. All study procedures were approved by the Institutional Review Board of the Technion–Israel Institute of Technology and were conducted in accordance with relevant ethical guidelines and regulations. Participants received compensation for the baseline scanning session and for completing the follow-up assessment.

### War-related exposure

To evaluate whether the primary connectivity–anxiety association was robust to adjustment for measured differences in war-related exposure during the study period, participants completed a war-related exposure questionnaire at follow-up (T2) assessing reserve service, war-related harm, relocation, frequency of exposure to alarms, and socioeconomic deterioration. Each item was converted into a binary indicator coded as 0 (not exposed) or 1 (exposed), according to the criteria reported in **Supplementary Table S3**. Reserve-service exposure was coded as present when participants reported that they or someone close to them was currently serving in the military reserves. War-related harm was coded as present when participants reported harm to themselves or an immediate family member. Relocation was coded as present when participants reported that they had not continued living in the same place of residence. Alarm exposure was coded as present when participants reported alarms in their area of residence once per day or more. Socioeconomic deterioration was coded as present when participants reported any degree of deterioration in their socioeconomic situation; the response “not at all” was coded as not exposed.

A security-exposure index was calculated as the unweighted sum of reserve-service exposure, war-related harm, relocation, and alarm exposure once per day or more (range, 0–4), with higher scores indicating greater cumulative exposure to security-related stressors. A full exposure index was calculated by adding the socioeconomic-deterioration indicator to the four security-related indicators (range, 0–5). Spearman correlations were used to examine associations of the security- and full-exposure indices, as well as the socioeconomic-deterioration indicator separately, with baseline right nucleus accumbens–right amygdala functional connectivity and subsequent anxiety symptoms. These analyses were secondary and exploratory, and the reported P values were two-sided and uncorrected. Finally, we evaluated the stability of the primary connectivity–anxiety association in regression models that incrementally adjusted for the security-exposure index, the full exposure index, and the security-exposure index and socioeconomic-deterioration indicator entered as separate predictors (**Supplementary Tables S4–S5** and **Supplementary Figs. S6–S7**).

### Baseline resting-state fMRI and in-scanner mood assessment

At baseline (T1), participants completed a 6 min 45 s resting-state fMRI scan. During the scan, participants fixated on a white cross presented on a black background and simultaneously also provided repeated ratings of their current mood. Each mood rating epoch lasted up to 7.5 s, comprising a 3.5-s question-presentation period followed by a 4-s rating period, and then followed by a 15-s resting interval. A total of 18 mood ratings were obtained during the scan. Mood ratings were obtained using a continuous visual analogue scale (VAS) in response to the question “How do you feel right now?” The scale was anchored by “unhappy” and “happy.” Participants moved a cursor along the scale using a continuous button press.

Because mood ratings were interleaved with the resting-state acquisition, rating epochs were modeled separately and regressed out during functional connectivity estimation to minimize the contribution of explicit mood reporting to estimates of intrinsic functional connectivity^16–18^.

### Psychological assessments

Depressive symptoms were assessed at baseline (T1) and follow-up (T2) using the 20-item Center for Epidemiologic Studies Depression Scale (CES-D)^19^. CES-D scores were analyzed primarily as a continuous measure of depressive symptom severity. The conventional threshold of CES-D ≥16 was used to characterize elevated depressive symptoms for descriptive and visualization purposes and was not used as the primary analytical outcome.

At follow-up (T2), anxiety symptoms were assessed using the Generalized Anxiety Disorder-7 (GAD-7)^20^, perceived stress using the Perceived Stress Scale (PSS)^21^, and post-traumatic stress symptoms using the PTSD Checklist for DSM-5 (PCL-5)^22^. Interoceptive awareness was additionally assessed using the Multidimensional Assessment of Interoceptive Awareness, Version 2 (MAIA-2)^23^. Questionnaire scores were calculated according to standard scoring procedures and analyzed as continuous measures.

The four psychological outcomes included in the prospective ROI-to-ROI functional connectivity analysis were follow-up CES-D, GAD-7, PSS, and PCL-5 scores. Anxiety symptoms were not assessed at baseline; accordingly, analyses involving GAD-7 quantify associations with subsequent anxiety symptom severity rather than within-participant change in anxiety from baseline.

### MRI acquisition

MRI data were acquired using a 3-T Siemens Prisma scanner equipped with a 64-channel head coil. Functional images were acquired using a gradient echo-planar imaging (EPI) sequence with the following parameters: repetition time (TR) = 2,500 ms, echo time (TE) = 22 ms, flip angle = 80°, field of view = 204 mm, acquisition matrix = 82 × 82, voxel size = 2.5 mm isotropic, 60 axial slices, and no interslice gap.

High-resolution T1-weighted anatomical images were acquired before the resting-state scan using a magnetization-prepared rapid gradient-echo (MPRAGE) sequence with the following parameters: TR = 3,150 ms, TE = 1.37 ms, flip angle = 8°, field of view = 260 mm, acquisition matrix = 160 × 160, and slice thickness ≈ 1 mm. During the anatomical scan, participants viewed a brief neutral documentary about the Steller’s sea eagle.

### fMRI preprocessing

Functional MRI data were preprocessed using the CONN functional connectivity toolbox v22a and SPM12^16^. Preprocessing included functional realignment and unwarping, identification of potentially artifactual volumes using the Artifact Detection Tools (ART), normalization to Montreal Neurological Institute (MNI) standard space, tissue segmentation, and spatial smoothing using an 8-mm full-width at half-maximum Gaussian kernel.

Volumes were identified as potential outliers when framewise displacement exceeded 0.9 mm or global signal intensity exceeded 5 SD from the mean. Denoising included anatomical CompCor components derived from white matter and cerebrospinal fluid signals, estimated motion parameters and associated regressors, identified outlier volumes, session effects, and linear temporal trends. The residual BOLD time series were band-pass filtered between 0.008 and 0.09 Hz.

### Regions of interest and functional connectivity estimation

The primary neuroimaging analysis focused on nine corticolimbic and reward-related regions of interest (ROIs) that were predefined in the original cohort study^15^ based on their established roles in affective, salience, threat and reward processing^24–26^. These comprised the bilateral insula, anterior cingulate cortex (ACC), medial prefrontal cortex (mPFC), posterior cingulate cortex (PCC), bilateral nucleus accumbens (NAcc), and bilateral amygdala. ROIs were anatomically defined using the Harvard–Oxford Atlas as implemented in the CONN toolbox.

For each participant, mean denoised BOLD time series were extracted from each ROI. Pairwise Pearson correlations were calculated between ROI time series and transformed using Fisher’s r-to-z transformation to obtain participant-level functional connectivity estimates. The nine predefined ROIs yielded 36 unique pairwise functional connections. Only connectivity estimates obtained at baseline (T1) were used in the prospective analyses.

### Statistical analyses

All statistical analyses were conducted in Python. Unless otherwise specified, statistical tests were two-sided, and continuous variables are reported as mean ± SD.

#### Prospective analysis of ROI-to-ROI connectivity and psychological outcomes

To examine whether baseline intrinsic functional connectivity was prospectively associated with subsequent psychological outcomes, each of the 36 baseline ROI-to-ROI functional connectivity estimates was tested separately for association with depressive symptoms (CES-D), anxiety symptoms (GAD-7), perceived stress (PSS), and post-traumatic stress symptoms (PCL-5) assessed at follow-up (T2).

Associations were quantified using two-sided Spearman rank correlations (ρ). For each psychological outcome, Benjamini–Hochberg false discovery rate (FDR) correction was applied across the 36 predefined functional connections. Thus, the multiple-comparison family comprised 36 connectivity tests for each outcome. FDR-adjusted q values < 0.05 were considered statistically significant. Where reported, 95% confidence intervals for Spearman correlation coefficients were estimated using 10,000 bootstrap resamples.

#### Multivariable analysis of subsequent anxiety symptoms

To determine whether the association between baseline right nucleus accumbens–right amygdala connectivity and subsequent anxiety symptoms remained evident after accounting for baseline depressive symptom severity and demographic characteristics, we fitted a multivariable linear regression model with follow-up GAD-7 score as the dependent variable. Baseline right nucleus accumbens–right amygdala functional connectivity was entered as the predictor of interest, with baseline CES-D score, age, and sex included as covariates. Sex was dummy-coded as 0 for male and 1 for female. For each predictor, we report the unstandardized regression coefficient (B), standardized coefficient (β), 95% confidence interval, and two-sided P value. Overall model fit was summarized using R², adjusted R², and the model F statistic.

#### In-scanner mood analyses

Repeated mood ratings obtained during the baseline scan were used to examine whether baseline connectivity was associated with participants’ concurrent affective state or within-scan mood dynamics. Six mood measures were derived: the first, mean, and final mood ratings; mood variability, quantified as the standard deviation of the ratings; overall mood change, calculated as the final rating minus the first rating (Δ mood); and mood slope. Individual mood slopes were estimated by fitting a linear regression across each participant’s repeated mood ratings, with more negative slopes indicating greater downward mood drift during the scan.

Associations between baseline right nucleus accumbens–right amygdala functional connectivity and the six in-scanner mood measures were assessed using two-sided Spearman rank correlations. Benjamini–Hochberg FDR correction was applied across the six measures, with q values < 0.05 considered statistically significant.

#### Sensitivity analyses

To control multiplicity across the full set of 144 connectivity–outcome associations, we performed a global permutation analysis using 100,000 permutations. In each permutation, a common permutation of participant labels was applied jointly to the four psychological outcomes relative to the connectivity data, thereby preserving the dependence structure among the outcomes. Empirical uncorrected P values for each association were calculated from the corresponding test-specific permutation distribution using the plus-one correction. Benjamini–Hochberg FDR correction was then applied across all 144 empirical P values.

In addition, the maximum absolute Spearman correlation across all 36 connections and four outcomes was retained in each permutation to construct a global maximum-statistic null distribution. Family-wise error-corrected P values were calculated by comparing each observed absolute correlation with this distribution. For the anxiety outcome, an outcome-specific maximum-statistic correction was also performed across the 36 predefined functional connections. In each permutation, the maximum absolute Spearman correlation across the 36 connectivity–anxiety associations was retained, and the observed correlation was compared with this outcome-specific null distribution to obtain a family-wise error-corrected P value. Statistical significance was defined as a global FDR-adjusted q value or family-wise error-corrected P value < 0.05.

## Data and code availability

The data supporting the findings of this study and the code used to perform the analyses and generate the figures will be made available through the Open Science Framework (OSF) upon publication, subject to applicable ethical and privacy restrictions.

