## Supplementary Material for "Pre-stress functional brain connectivity is associated with subsequent anxiety during war-related stress"

[1] The Azrieli Faculty of Medicine, Bar-Ilan University, Safed, Israel
[2] The Leslie and Susan Gonda Multidisciplinary Brain Research Center, Bar-Ilan University, Ramat Gan, Israel
* These authors contributed equally to this work

### Supplementary

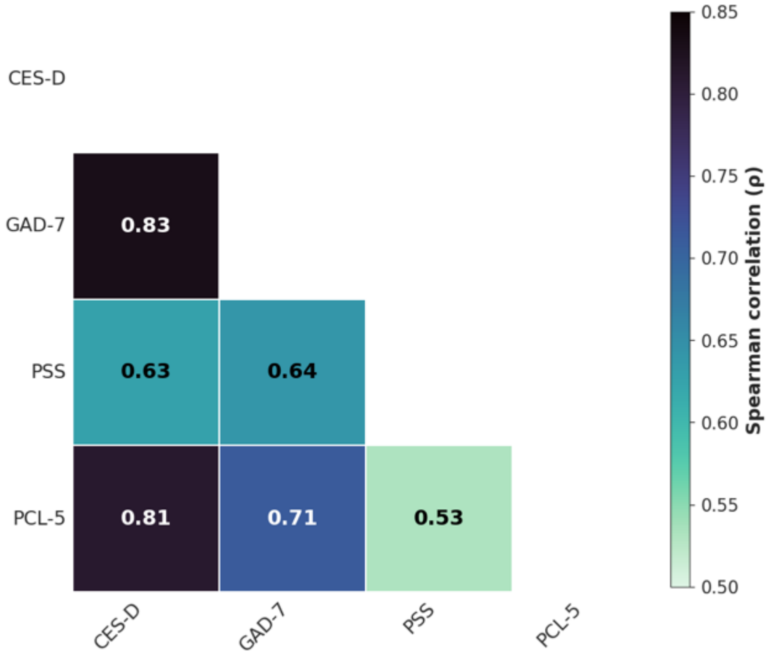


**Figure S1. Intercorrelations among follow-up clinical measures.** Lower triangular heatmap showing pairwise Spearman correlation coefficients (ρ) among follow-up depressive symptoms (CES-D), anxiety symptoms (GAD-7), perceived stress (PSS), and post-traumatic stress symptoms (PCL-5) in the fMRI cohort ($N = 31$). Darker colors indicate stronger positive correlations. Numbers within each cell indicate Spearman correlation coefficients (ρ). All pairwise correlations remained significant after Benjamini–Hochberg false discovery rate (FDR) correction across the six pairwise tests.

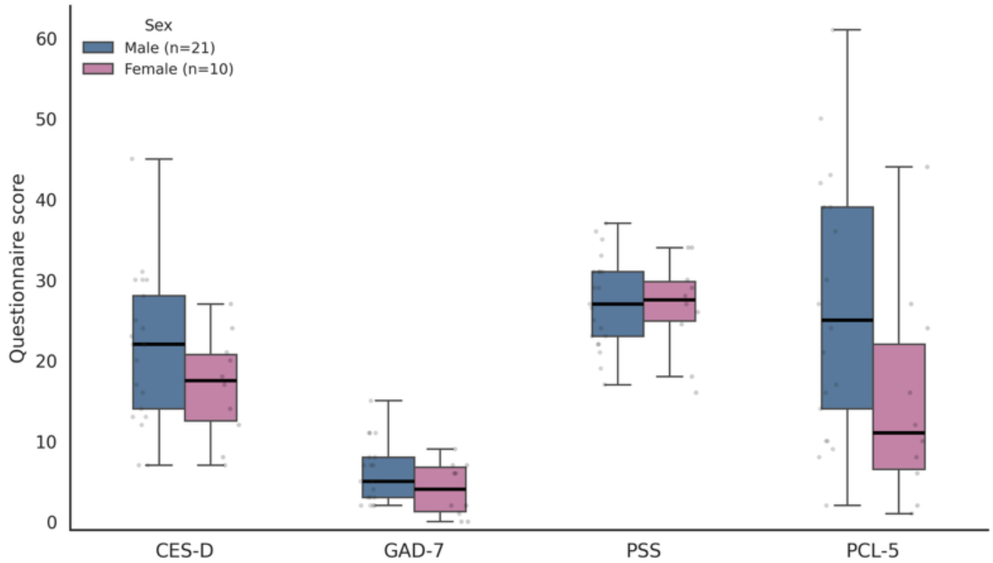


**Figure S2. Clinical questionnaire scores by sex in the fMRI cohort.** Distribution of follow-up CES-D, GAD-7, PSS, and PCL-5 scores in male ($n$ = 21) and female ($n$ = 10) participants from the fMRI cohort ($N$ = 31). Boxes represent the interquartile range (IQR), center lines indicate the median, whiskers extend to the most extreme values within 1.5 × IQR, and points represent individual participants. Sex differences were assessed using two-sided Welch's $t$-test with Benjamini–Hochberg false discovery rate (FDR) correction applied across the four questionnaire measures. No significant sex differences were observed after FDR correction.

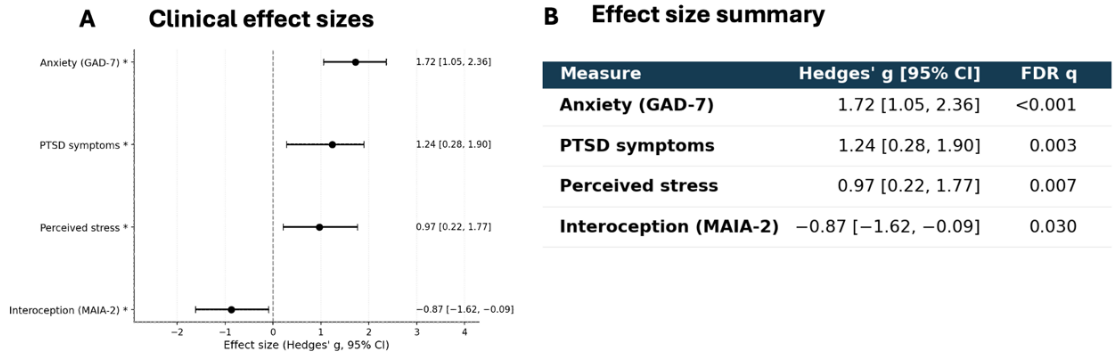


| Measure | Hedges' g [95% CI] | FDR q |
|---|---|---|
| **Anxiety (GAD-7)** | 1.72 [1.05, 2.36] | <0.001 |
| **PTSD symptoms** | 1.24 [0.28, 1.90] | 0.003 |
| **Perceived stress** | 0.97 [0.22, 1.77] | 0.007 |
| **Interoception (MAIA-2)** | −0.87 [−1.62, −0.09] | 0.030 |

**Figure S3. Clinical profile of participants with elevated depressive symptoms at follow-up.** (**A**) Forest plot showing Hedges' g and 95% bias-corrected and accelerated (BCa) bootstrap confidence intervals for follow-up clinical measures comparing participants with elevated depressive symptoms **(**CES-D ≥ 16; n = 21**)** and participants below the clinical threshold **(**CES-D < 16; n = 10**).** Positive effect sizes indicate higher scores in the elevated CES-D group. Asterisks indicate associations that remained significant after Benjamini–Hochberg false discovery rate (FDR) correction. (**B**)

Summary of effect sizes, 95% confidence intervals, and FDR-adjusted q values across the four clinical measures.

| Predictor | Coefficient (B) | Standardized β | 95% CI | P value |
|---|---|---|---|---|
| **Right NAcc–amygdala FC** | **−9.50** | **−0.53** | **−15.14 to −3.86** | **0.002** |
| Baseline CES-D | 0.13 | 0.28 | −0.02 to 0.28 | 0.094 |
| Age | 0.10 | 0.08 | −0.28 to 0.48 | 0.602 |
| Sex | −0.22 | −0.06 | −2.81 to 2.37 | 0.860 |
| ***Model statistics*** | | | | |
| N = 31 | $R^2$ = 0.443 | Adjusted $R^2$ = 0.357 | F(4, 26) = 5.16 | Model P = 0.003 |

**Table S1. Multivariable regression model of follow-up anxiety symptoms.** Note. **N = 31**. Follow-up GAD-7 score was entered as the dependent variable. B denotes the unstandardized regression coefficient; β denotes the standardized regression coefficient; CI denotes 95% confidence interval; FC denotes functional connectivity; NAcc denotes nucleus accumbens; CES-D denotes the Center for Epidemiologic Studies Depression Scale. Sex was coded as 0 = male and 1 = female. The overall model fit was statistically significant (F(4, 26) = 5.16, P = 0.003, $R^2$ = 0.443, adjusted $R^2$ = 0.357).

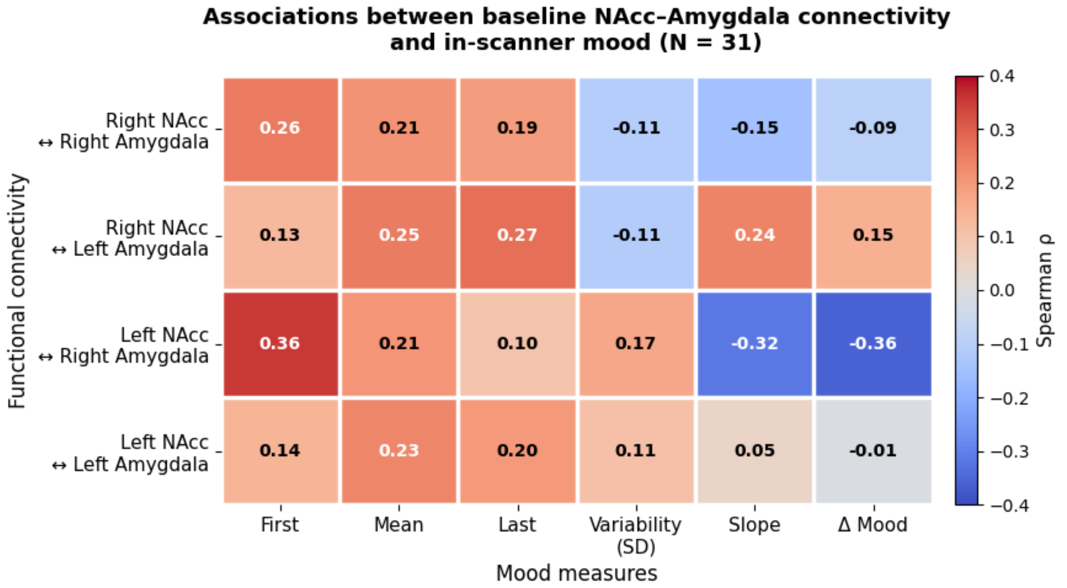


**Figure S4. Associations between baseline NAcc–amygdala connectivity and in-scanner mood dynamics (N = 31).** Heatmap showing Spearman correlation coefficients between baseline resting-state functional connectivity of four nucleus accumbens (NAcc)–amygdala functional connections and six in-scanner mood measures (first, mean, and last mood ratings, mood variability, mood slope, and overall mood change) in 31 participants. Cell values indicate Spearman's correlation coefficients (ρ), with warmer colors representing positive associations and cooler colors representing negative associations. P values were adjusted for multiple comparisons using the Benjamini–Hochberg false discovery rate (FDR) procedure. No associations remained significant after FDR correction.

| Analysis | Spearman ρ | Standardized β | B | 95% CI / range | P or q value |
|---|---|---|---|---|---|
| ***Correlation-based robustness analyses*** | | | | | |
| **Primary Spearman correlation** | **−0.619** | — | — | — | **$2.02 \times 10^{-4}$** |
| Leave-one-out analysis | Mean: −0.619 | — | — | −0.668 to −0.580 | — |
| Bootstrap (10,000 resamples) | Median: −0.618 | — | — | [−0.807, −0.314][b] | — |
| Permutation test (100,000 permutations) | Observed: −0.619 | — | — | — | $3.20 \times 10^{-4}$[b] |
| Global BH FDR correction (144 associations) | Observed: −0.619 | — | — | — | q = 0.046[d] |
| Global max-statistic correction (144 associations) | Observed: −0.619 | — | — | — | P(FWE) = 0.032[a] |
| ***Regression sensitivity analyses*** | | | | | |
| Reduced regression: FC + baseline CES-D | — | −0.534 | −9.49 | [−14.79, −4.19][c] | 0.0010 |
| Full regression: FC + CES-D + age + sex | — | −0.534 | −9.50 | [−15.14, −3.86][c] | 0.0019 |
| Full regression with HC3 robust SE | — | — | −9.50 | [−16.03, −2.96][c] | 0.0044 |

**Table S2. Sensitivity and robustness analyses of the primary right nucleus accumbens–right amygdala functional connectivity–anxiety association.** Note. **N = 31** for all analyses. FC = functional connectivity; NAcc = nucleus accumbens; GAD-7 = Generalized Anxiety Disorder-7; CES-D = Center for Epidemiologic Studies Depression Scale; CI = confidence interval; HC3 = heteroskedasticity-consistent (HC3) standard errors; FDR = false discovery rate; FWE = family-wise error. All regression models used subsequent GAD-7 score as the dependent variable. Sex was dummy-coded as 0 = male and 1 = female. Standardized β and unstandardized B are reported in separate columns; confidence intervals for the regression coefficients refer to unstandardized B, not standardized β. The reduced model included right NAcc–right amygdala FC and baseline CES-D; the full model additionally included age and sex. There was no evidence of problematic multicollinearity in the full model (all predictor VIFs ≤ 1.28; FC VIF = 1.11). [a] 95% percentile bootstrap CI for ρ. [b] Empirical two-sided permutation P value calculated using the plus-one correction. [c] 95% CI for unstandardized B. [d] Benjamini–Hochberg FDR-adjusted q value across all 144 connectivity–outcome associations. [e] Permutation-based maximum-statistic P value corrected across all 144 connectivity–outcome associations; participant labels for the four outcomes were permuted jointly to preserve the dependence structure among outcomes.

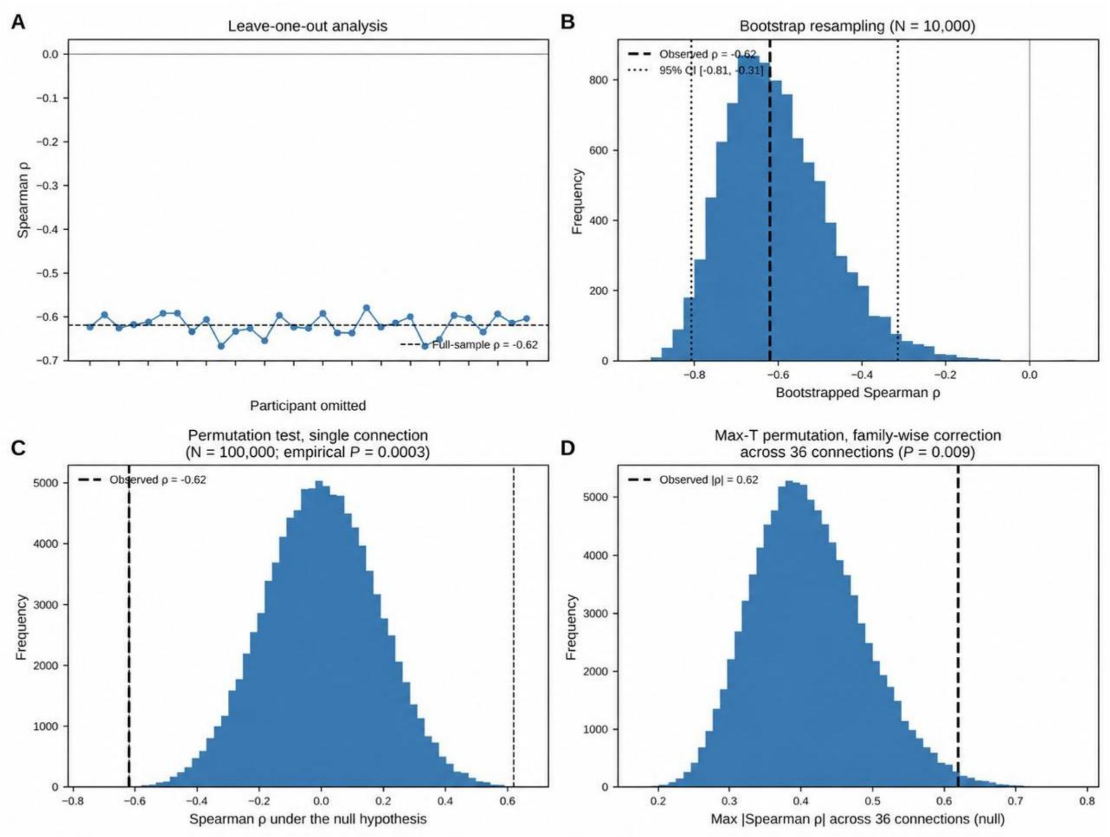


**Figure S5. Robustness of the primary right nucleus accumbens–right amygdala functional connectivity–anxiety association. (A)** Leave-one-out analysis of the association between baseline right nucleus accumbens–right amygdala functional connectivity and subsequent anxiety symptoms (GAD-7). Each point represents the Spearman correlation coefficient (ρ) recalculated after omitting one participant at a time (N = 30 per iteration). The dashed line indicates the full-sample correlation coefficient (ρ = −0.62). The correlation remained stable across all 31 iterations (ρ range, −0.67 to −0.58), indicating that the association was not driven by any single participant. (B) Bootstrap distribution of the primary functional connectivity–anxiety association. The histogram shows the distribution of Spearman correlation coefficients obtained from 10,000 bootstrap resamples of the original dataset (N = 31). The dashed line indicates the observed correlation coefficient, and the dotted lines indicate the 95% percentile bootstrap confidence interval (95% CI, −0.81 to −0.31), which excluded zero. **(C)** Permutation null distribution of the primary functional connectivity–anxiety association. The histogram shows the distribution of Spearman correlation coefficients obtained from 100,000 permutations in which follow-up anxiety scores were randomly reassigned across participants. The dashed line indicates the observed correlation coefficient (ρ = −0.62), which exceeded the magnitude of all but 30 of the 100,000 permuted values (empirical two-sided P = 0.0003), yielding a result consistent with the analytic test**.** **(D)** Outcome-specific maximum-statistic permutation null distribution controlling the family-wise error rate across the 36 baseline functional connections examined in the ROI-to-ROI connectivity screen (**Fig. 2**). For each of 100,000 permutations, follow-up anxiety scores were randomly reassigned once and correlated with all 36 connections simultaneously; the maximum absolute correlation coefficient across the 36 connections was retained to construct the null distribution. The dashed line indicates the observed absolute correlation coefficient for the right nucleus accumbens–right amygdala connection, which remained significant after family-wise error correction (P_FWE = 0.009).

| Exposure indicator | Survey item (translated from Hebrew) | Coded as exposed (= 1) when... | N exposed / 31 (%) |
|---|---|---|---|
| Reserve service | Are you, or someone close to you, currently serving in the military reserves? | Response indicated any current reserve service (any option other than "not serving") | 15 (48.4) |
| War-related harm | Have you or your immediate family members been harmed during the war? (multiple selections allowed) | Response indicated any form of harm (any option other than "not harmed") | 2 (6.5) |
| Relocation | Did you continue living in the same place of residence during this period? | Response indicated a change in place of residence | 9 (29.0) |
| Frequent alarms | How frequently are there alarms/sirens in your area of residence? | Response indicated alarms once per day or more | 19 (61.3) |
| Socioeconomic deterioration | Has your socioeconomic situation worsened in the past month? (e.g., business closure, unpaid leave) | Response indicated any degree of deterioration (codes 2–5; code 1 = not at all) | 11 (35.5) |

**Table S3. War-related exposure items: wording, coding, and prevalence in the fMRI cohort. N = 31**. Items are English translations of a Hebrew war-related exposure questionnaire administered at follow-up (T2). Original response options were categorical and not necessarily equal-interval; responses were collapsed into binary indicators coded as exposed or not exposed according to the criteria shown. The five binary indicators were combined into two unweighted cumulative indices: a security-exposure index comprising reserve service, war-related harm, relocation, and alarm exposure once per day or more (range, 0–4), and a full exposure index comprising the four security-related indicators plus socioeconomic deterioration (range, 0–5). Higher scores indicate greater cumulative exposure**.** The indices were used in the exposure-adjusted analyses reported in **Tables S4–S5**.

| Variable 1 | Variable 2 | N | Spearman $\rho$ | P |
|---|---|---|---|---|
| **War-related exposure analyses** | | | | |
| Security-exposure index | Right NAcc–right amygdala FC | 31 | −0.112 | .5480 |
| Security-exposure index | Subsequent anxiety symptoms (GAD-7) | 31 | 0.377 | .0365 |
| Full exposure index | Right NAcc–right amygdala FC | 31 | −0.144 | .4409 |
| Full exposure index | Subsequent anxiety symptoms (GAD-7) | 31 | 0.444 | .0124 |
| Socioeconomic deterioration | Right NAcc–right amygdala FC | 31 | −0.219 | .2374 |
| Socioeconomic deterioration | Subsequent anxiety symptoms (GAD-7) | 31 | 0.422 | .0182 |
| **Primary connectivity association** | | | | |
| **Right NAcc–right amygdala FC** | Subsequent anxiety symptoms (GAD-7) | 31 | −0.619 | .0002 |

**Table S4. Associations between war-related exposure, baseline connectivity, and subsequent anxiety symptoms. N = 31** for all analyses. FC, functional connectivity; NAcc, nucleus accumbens; GAD-7, Generalized Anxiety Disorder-7. The security-exposure index comprises reserve service, war-related harm, relocation, and alarm exposure once per day or more (range, 0–4); the full exposure index additionally includes socioeconomic deterioration (range, 0–5; **Table S3**). Neither exposure index was associated with baseline right NAcc–right amygdala connectivity. In uncorrected secondary analyses, the security-exposure index, full exposure index, and socioeconomic-deterioration indicator each showed positive associations with subsequent anxiety symptoms. Socioeconomic deterioration was not associated with baseline connectivity. Exposure-adjusted regression models are reported in **Table S5**. All reported P values are two-sided and uncorrected.

| Model | B | HC3 SE | 95% CI lower | 95% CI upper | P | $R^2$ | Adjusted $R^2$ |
|---|---|---|---|---|---|---|---|
| FC + baseline CES-D | −9.493 | 3.227 | −16.103 | −2.883 | .0065 | .437 | .396 |
| + age and sex | −9.498 | 3.335 | −16.353 | −2.644 | .0085 | .443 | .357 |
| + security-exposure index | −9.284 | 2.658 | −14.758 | −3.810 | .0018 | .545 | .454 |
| + full exposure index | −8.569 | 2.165 | −13.029 | −4.110 | .0006 | .623 | .547 |
| + security-exposure index and socioeconomic deterioration | −8.196 | 2.197 | −12.730 | −3.661 | .0010 | .640 | .550 |

**Table S5. Stability of the right NAcc–right amygdala connectivity coefficient across models adjusting for war-related exposure.** Note. **N = 31** for all models. FC, functional connectivity; NAcc, nucleus accumbens; CES-D, Center for Epidemiologic Studies Depression Scale; GAD-7, Generalized Anxiety Disorder-7; HC3, heteroskedasticity-consistent standard errors; CI, confidence interval. All models used subsequent GAD-7 score as the dependent variable and were estimated using HC3-robust standard errors and t-distribution-based inference, with residual degrees of freedom calculated as df = N − k, where k denotes the number of estimated model parameters. Models sequentially adjusted for baseline CES-D, age and sex; the security-exposure index (**Table S3**); the full exposure index, which additionally included socioeconomic deterioration; and the security-exposure index and socioeconomic-deterioration indicator entered as separate predictors. The right NAcc–right amygdala connectivity coefficient remained similar in magnitude across models (B range, −9.50 to −8.20) and remained statistically significant in all models (all $P < 0.009$), indicating that the association between baseline connectivity and subsequent anxiety remained evident after adjustment for the measured war-related exposure indicators.

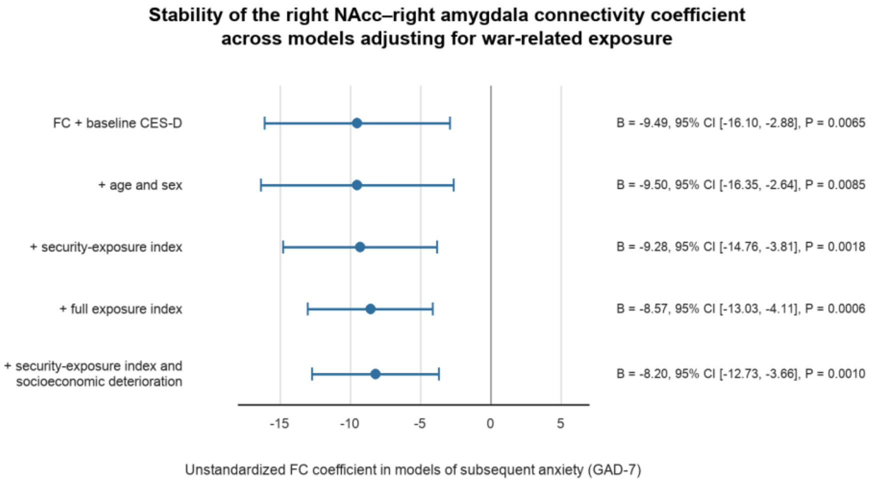


**Figure S6. Stability of the right NAcc–right amygdala connectivity coefficient across models adjusting for war-related exposure.** Forest plot of the unstandardized regression coefficient (B) for right NAcc–right amygdala functional connectivity in models of subsequent anxiety symptoms (GAD-7), estimated across five progressively adjusted model specifications using HC3-robust standard errors and t-distribution-based inference (**N = 31** in all models). Circles represent point estimates; error bars represent 95% confidence intervals; and the dashed vertical line marks zero. Models included a base model containing FC and baseline CES-D, followed by adjustment for age and sex; the security-exposure index; the full exposure index, which additionally included socioeconomic deterioration; and the security-exposure index and socioeconomic-deterioration indicator entered as separate predictors. The connectivity coefficient remained similar in magnitude, and its confidence interval excluded zero in every model, indicating that the association between baseline connectivity and subsequent anxiety remained evident after adjustment for the measured war-related exposure indicators. Full model statistics are reported in **Table S5**.

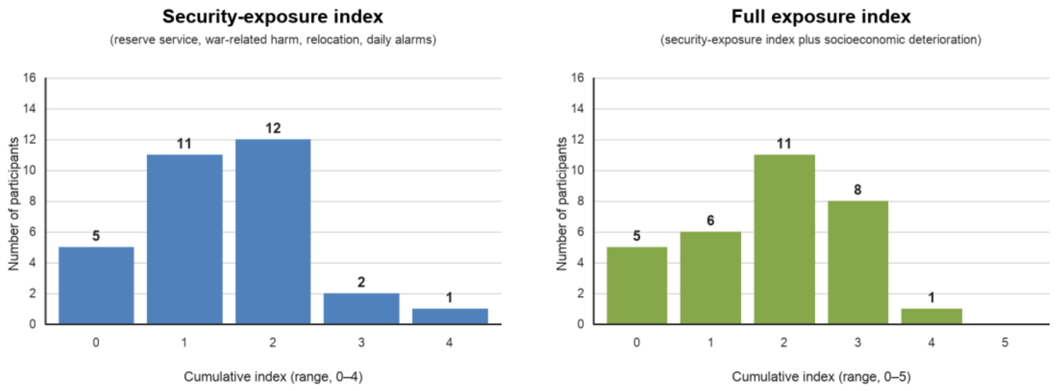


**Figure S7. Distribution of cumulative war-related exposure indices (N = 31).** Distribution of participant scores on the two cumulative war-related exposure indices constructed from the binary indicators described in **Table S3**. Left, security-exposure index, calculated as the sum of reserve service, war-related harm, relocation, and alarm exposure once per day or more (range, 0–4). Right, full exposure index, calculated as the security-exposure index plus socioeconomic deterioration (range, 0–5). Both indices varied across the cohort and were positively associated with subsequent anxiety symptoms in uncorrected secondary analyses (**Table S4**).
